# A whole nervous system atlas of glutamate receptors reveals distinct receptor roles in sensorimotor circuits

**DOI:** 10.1101/2023.04.18.537384

**Authors:** Cezar Borba, Matthew J. Kourakis, Yishen Miao, Bharath Guduri, Jianan Deng, William C. Smith

## Abstract

A goal of connectomics is to reveal the links between neural circuits and behavior. Larvae of the primitive chordate *Ciona* are well-suited to make contributions in this area. The small size of the *Ciona* larval nervous system (∼180 neurons) has facilitated the full characterization of a connectome. In addition, the larvae display a range of behaviors that are readily quantified in both normal and manipulated larvae. Moreover, the small number of neurons allows for a neuron-by-neuron characterization of attributes such as neurotransmitter use. We present here a nervous system-wide atlas of glutamate receptor expression. Included in the atlas are both ionotropic receptors (AMPA, NMDA, and Kainate), and metabotropic receptors. The expression of these receptors is presented in the context of known circuits driving behaviors such as phototaxis, mechanosensation, and looming shadow response. The expression of AMPA and NMDA receptors, in particular, helps to resolve the apparently paradoxical coproduction of GABA and glutamate by some photoreceptors. We find that the targets of these photoreceptors, midbrain relay neurons, primarily express NMDA receptors in the absence of AMPA receptors. This is in agreement with previous results indicating that GABA is the primary neurotransmitter from the photoreceptors evoking a behavioral response (swimming) through a disinhibition mechanism. We hypothesize that NMDA receptors have a modulatory effect in the relay neurons. Other findings reported here are more unexpected. For example, the targets of glutamatergic epidermal sensory neurons (ESNs) do not express any of the ionotropic receptors, yet the ESNs themselves express metabotropic receptors. Thus, we speculate that their production of glutamate may be for communication with neighboring ESNs, rather than to their interneuron targets.

## Introduction

The tadpole larva of the invertebrate chordate *Ciona* is a highly tractable model for sensorimotor circuit analyses. Not only does the *Ciona* larval central nervous system (CNS) contain only ∼180 neurons, it is one of the few animals for which a complete synaptic connectome has been described (Ryan et al., 2016). Moreover, numerous studies have highlighted the conservation between the *Ciona* larval CNS and those of vertebrates [reviewed in (Hudson, 2016)]. At the anatomical level, the *Ciona* CNS is subdivided into domains showing homology to the vertebrate forebrain, midbrain, hindbrain, midbrain-hindbrain boundary (MHB), and spinal cord (Figure 1a). These homologies are evident in *Ciona’s* developmental mechanisms, gene expression, anatomy, and most recently in neuron classification and synaptic connectivity (Borba et al., 2021; Hudson, 2016; Ryan et al., 2017; Wada et al., 1998). Early descriptions of larval tunicate nervous systems, often made before the above homologies were clear, led to the naming of these anatomical domains with names that do not reflect this homology (*e.g.*, *anterior sensory vesicle* or *anterior brain vesicle* for the *Ciona* forebrain homolog, and *visceral* or *motor ganglion* for the hindbrain homolog). For the sake of clarity, and to make *Ciona* neurobiology accessible to a broader readership, we will henceforth refer to the *Ciona* CNS anatomical domains according to their vertebrate homologs.

**Figure 1.**
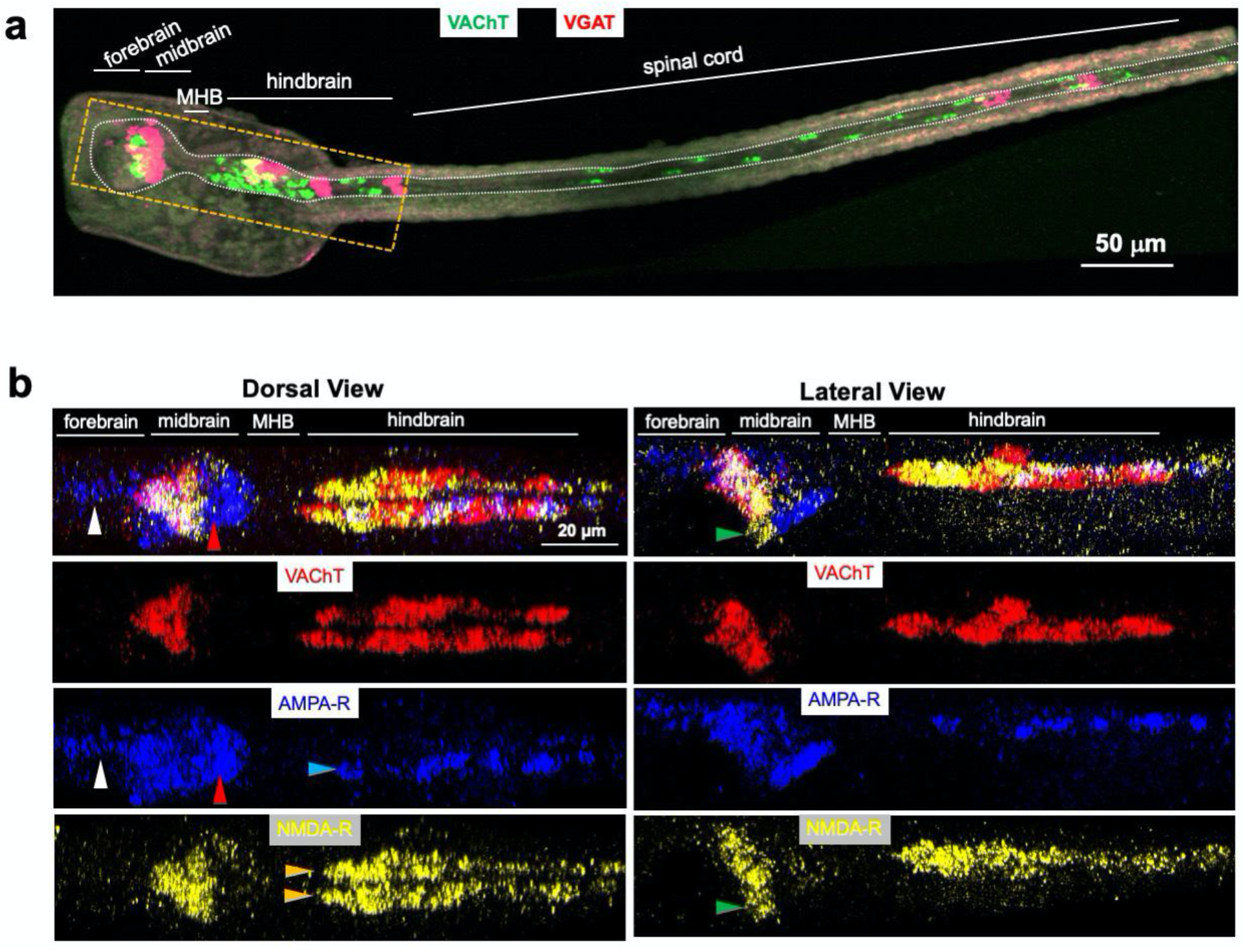
*In situ* hybridization of *C. robusta* larvae. **a**. *In situ* hybridization of *C. robusta* larva for VAChT and VGAT. The major subdivisions of the central nervous system are labeled according to their vertebrate orthologs. The white line outlines the central nervous system, and the orange box outlines the approximate brain regions shown below in panel b. White arrows indicate midtail neurons. **b.** *In situ* hybridization of *C. robusta* larva for VAChT, AMPA-R and NMDA-R. Top panels show dorsal and lateral composite views of the three expression patterns. The three lower right and left panels show the individual images that comprise the composite. White and red arrowheads indicate AMPA-R^+^/NMDA-R^-^ forebrain and posterior midbrain neurons, respectively. Green arrowheads indicate NMDA-R^+^/AMPA-R^-^ neurons. All images are maximum intensity Z-projection images from confocal stacks. Anterior is to the left for all panels.

The *Ciona* connectome dataset has been instrumental in identifying neural circuits that drive a number of larval behaviors, including negative phototaxis, negative gravitaxis, a touch response, and a looming shadow response (Bostwick et al., 2020; Kourakis et al., 2019; Ryan et al., 2018, 2016). The connectome, however, which was derived by serial-section electron microscopy, provides only a “bare-bones” view of the neural circuitry. A fuller understanding of the logic of neural circuits requires that “flesh” be added in the form of the properties of the constituent neurons (*e.g.*, neurotransmitter use and neurotransmitter receptor expression). A detailed picture of the neurotransmitter use in the *Ciona* larval CNS is emerging through analysis of *in situ* hybridization patterns for markers of small-molecule neurotransmitters [*e.g.*, vesicular acetylcholine transporter (VAChT) for cholinergic neurons, vesicular GABA transporter (VGAT) for GABAergic/glycinergic neurons, tyrosine hydroxylase (TH) for catecholaminergic neurons, and tryptophan hydroxylase (TPH) for serotonergic neurons] (Kourakis et al., 2019; Moret et al., 2005; Pennati et al., 2007). The low number of larval neurons, and the largely stereotyped expression patterns, facilitate the mapping of neurotransmitter use to the connectome (Kourakis et al., 2019). For example, all vesicular glutamate transporter (VGLUT)-positive neurons in the *Ciona* larva are sensory (*i.e.*, there are no glutamatergic interneurons or efferents). The list of glutamatergic neurons includes the photoreceptors, the gravity-sensitive antennae cells, and the peripheral epidermal sensory neurons (Horie et al., 2008a; Kourakis et al., 2019). TH expression is limited to the photoreceptor-like coronet cells, and these neurons are presumed to be dopaminergic - although the *Ciona* genome appears to lack a definitive dopamine receptor (Lemaire et al., 2021; Moret et al., 2005). Interneurons of the CNS are either VGAT^+^, VAChT^+^, or express none of the markers of small molecule neurotransmitters, and are therefore likely peptidergic (Hamada et al., 2011; Kourakis et al., 2019). In the *Ciona* midbrain, VAChT and VGAT are expressed in distinct, non-intermingled domains, with the VAChT domain anterior to the VGAT domain, while in the hindbrain VAChT expression dominates with VGAT expression limited to six *ascending motor ganglion* peripheral interneurons (AMGs) in the dorsal midbrain, and four *ascending contralateral inhibitory neurons* (ACINs) (Kourakis et al., 2019).

In order to better understand the role of the glutamate system in *Ciona* behavior and neural circuitry, and to complement our knowledge of neurotransmitter distribution, we undertook a comprehensive examination of the expression patterns of the glutamatergic ionotropic receptors (AMPA, NMDA, and kainate) and metabotropic receptors (mGluR) at the larval stage using hybridization chain reaction *in situ* (Choi et al., 2018). While *Ciona* and related animals (the tunicates) are the closest extant relatives of the vertebrates, they diverged in a number of important ways from the vertebrates. Significantly, the tunicates split from the vertebrates before two whole-genome duplications occurred in the vertebrate lineage (Dehal and Boore, 2005). As a result, tunicates have smaller genomes when compared to vertebrates, and in most cases have fewer members of gene families. This relationship is evident in the glutamate receptors: the *Ciona* genome encodes single copies of the AMPA and kainate receptors, as well as single copies of each NMDA receptor subunit (Okamura et al., 2005), and three genes putatively encoding mGlu receptors (Kamesh et al., 2008). Thus, the relative genomic simplicity of *Ciona* greatly simplifies the task of generating a comprehensive view of the expression of the glutamate receptors, while the relatively simple behaviors of *Ciona* larvae, and the circuits driving them, aid in the assessment of the roles of the glutamatergic receptors.

## Material and Methods

### Animals

Adult *Ciona robusta* [also known as *Ciona intestinalis* type A] were collected at the Santa Barbara Harbor. Gametes were dissected from adults and crossed *in vitro* to generate larvae. All embryos and larvae were cultured at 18°C.

### *In situ* hybridization and image collection

Whole mount fluorescent *in situ* hybridization of larval *C. robusta* were performed using the hybridization chain reaction (HCR) method (v. 3.0, Molecular Instruments; Los Angeles, CA, United States), as previously described (Kourakis et al., 2019). Complementary RNA probe sets were designed to coding regions for the following *Ciona* genes (unique gene identifiers provided in parentheses): AMPA receptor (XM_018817034.1), NMDA receptor (XM_018816819.1), Kainate receptor (XM_026833998.1), metabotropic glutamate receptor 123 (XM_009859697.3), metabotropic glutamate receptor 478 (XM_018816381.1), VGAT (NM_001032573.1), and VAChT (NM_001032789.1). Larvae for *in situ* hybridization were dechorionated at mid-tailbud stage using sodium thioglycolate/protease E or 0.1% trypsin, so that left-right asymmetric properties of the CNS would not be disrupted, as described previously (Kourakis et al., 2021). Labeled animals were imaged on an Olympus Fluoview 1000 confocal microscope; post-image analysis used Imaris v6.4.0.0 or ImarisViewer v9.5.1 as well as Fiji (ImageJ) v. 2.0.0-rc-69/1.52p.

### Mapping of *in situ* hybridization patterns to the connectome

The TIFF image stack for *in situ* HCR expression was converted into voxels through a custom MATLAB and C# script. In MATLAB, the expression value of a given pixel in a z-plane of the TIFF stack is converted into an 8-bit integer (0-255). Each z-plane in the stack is thus represented by a CSV table where the row- and column-coordinate of the integer expression value are the xy-coordinates of the corresponding pixel in that plane. Then, with the C# script, each voxel was individually loaded into a custom Unity project for each TIFF stack, where the volume of a voxel is determined by the scale of the pixel and the distance between the z-planes. As the voxels are loaded, their meshes are merged and saved as a Unity asset. The final mesh is a 3D construct object of the *in situ* HCR expression result. The cell shape reconstruction from the connectome data (Ryan et al., 2016) were loaded into the same scene as expression objects for alignment. In order to load the cell shapes, the original data was loaded into the program Reconstruct, where it was exported as a scene. The scene was loaded into Blender where each individual neuron was exported as a .DXF file for loading into Unity. This consisted of two datasets: a “low-resolution” that contained all the data for the brain vesicle, and a “high-resolution” which contained all the data for the motor ganglion. Since VGAT expression is well characterized in both the brain vesicle and the motor ganglion (Kourakis et al., 2019), the expression objects were rotated and overlaid with the cell shapes manually using the following criteria in relation to VGAT expression: in the brain vesicle, the dorsal cap marks the Eminen cells and the two patches, a smaller posterior one and a larger anterior one, on the right mark the two photoreceptor groups, PR-I (only pr-9 and pr-10) and PR-II, respectively; in the motor ganglion, the dorsal patch of VGAT marks the AMG (all except AMG-5). After this alignment is done, expression objects with no consistent landmarks, such as VAChT, are brought into view. Using common structures and overlaps across different *in situs*, all the expression objects were overlapped in reference to each other. The custom software used here is available from https://github.com/CionaLab/agr.

### Analysis of mapped expression patterns

To align coordinates used in mapping, a Unity program using a custom C# script first loads the expression data voxel-by-voxel. The program checks if a voxel collides with the mesh of a cell shape. If there is a collision, the expression volume of that voxel is assigned to that neuron. The total expression percent is then determined by taking the total assigned expression volume and dividing by the total cell volume. The total expression percent of the neuron was then normalized to relative expression by dividing the value by the highest total expressing neuron for that specific *in situ*. The neuron was predicted to be positive for a particular transcript if the total expression percent was at least 5.5%. The threshold value was determined by first analyzing the hindbrain (also known as the motor ganglion), as the expression of VACHT, VGAT and AMPA-R has been described before (Kourakis et al., 2021, 2019). In particular, VACHT is expressed in AMG5, the six MGINs and all ten MNs, while VGAT is expressed in AMG1,2,3,4, 6 and 7. Finally, AMPA-R is expressed in the left motor neurons. Using hindbrain expression values guides, the analysis was performed with a variety of parameters until determining a threshold that aligns with this “ground truth”. These values were used to then determine fore- and midbrain expression.

### Behavioral Assays

All larvae were between 25 and 28 hours post fertilization (hpf) (18°C). Larval swimming behaviors were recorded in seawater using 10 cm agarose-coated petri dishes to reduce sticking. Image series were collected using a Hamamatsu Orca-ER camera fitted on a Navitar 7000 macro zoom lens. Programmable 700 and 505 nm LED lamps (Mightex) mounted above the petri dishes were used for dimming response assays as described previously (Borba et al., 2021; Bostwick et al., 2020; Kourakis et al., 2019). The dim response movies were recorded at 10 frames per second (fps). The larvae were recorded for 10 s at the initial intensity (3 mW/cm^2^) that was then dimmed (0.3 mW/cm^2^) while image capture continued for 1 min. Larvae were allowed to recover for 5 min before being assessed again. All light intensity readings were taken with an Extech Instruments light meter. For pharmacological experiments, MK801 (Tocris) was dissolved in seawater to a concentration of 500 μM, and the larvae were exposed to the drug for 10 minutes before being assessed.

## RESULTS

### Ionotropic glutamate receptors are expressed in broad, partially overlapping domains

*In situ* hybridization by HCR allows for the easy visualization of multiple fluorescently-labeled probes in a single sample (Choi et al., 2018). In present study, probes for ionotropic glutamate receptors (AMPA, NMDA, and kainate), as well as for the cholinergic marker VAChT, and the GABA/glycinergic marker VGAT, were tested in groups of three. For the NMDA receptor (NMDA-R), the *Ciona* homolog of the GluN1 subunit was used, as it is common to all NMDA-R complexes (Paoletti et al., 2013). Figure 1b shows 2D-projections of confocal images for the expression of VAChT with the AMPA and NMDA receptors in a representative 25 (hpf) larva, while Figure 2 shows the expression of VGAT with NMDA and kainate receptors in an identically-staged larva (see also SMovie 1 and 2). As seen in Figure 1b, the AMPA receptor (AMPA-R) expression domain in the fore- and midbrains is more extensive than the NMDA receptor. Expression of AMPA-R in the absence of NMDA-R was observed in the forebrain (white arrowhead), as well as the posterior midbrain (red arrowhead). We also observed a domain in the midbrain expressing NMDA-R, but not AMPA-R (green arrows in Figure 1b, lateral view). No expression of either NMDA-R or AMPA-R was observed in the MHB region.

**Figure 2.**
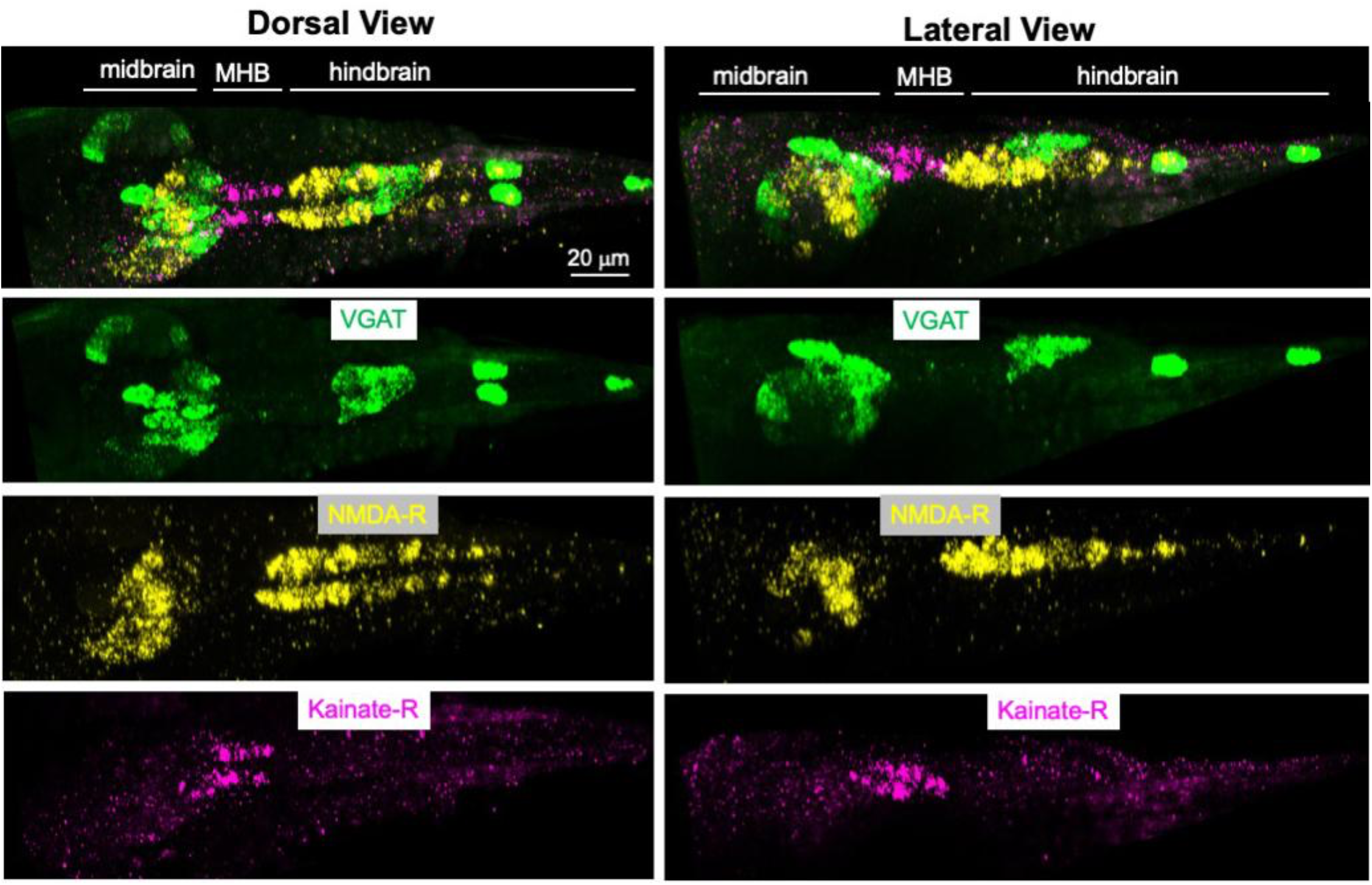
*In situ* hybridization of *C. robusta* larva for VGAT, NMDA-R, and Kainate-R. Top panels show dorsal and lateral composite views of the three expression patterns. The three lower right and left panels show the individual images that comprise the composite. White and red arrowheads indicate AMPA-R^+^/NMDA-R^-^ forebrain and posterior midbrain neurons, respectively. Green arrowheads indicate NMDA-R^+^/AMPA-R^-^ neurons. All images are Z-projection images from confocal stacks. Anterior is to the left for all panels. MHB, midbrain-hindbrain boundary.

In contrast to the fore- and midbrains, the expression of NMDA-R in the hindbrain was more extensive than that of the AMPA-R. With the exception of the seven dorsally-located AMG neurons (Ryan et al., 2018), the *Ciona* hindbrain is divided into distinct left and right sides. This is most conspicuous in the five cholinergic motor neurons found on each side which innervate tail muscles on the corresponding side (Ryan et al., 2016). However, this left/right symmetry is also present in the interneurons of the ventral hindbrain. Despite the symmetry of the ventral hindbrain at the level of neurons, we recently reported that AMPA-R is expressed only on the left side [(Kourakis et al., 2021) and blue arrowhead Figure 1b, dorsal view]. By contrast, we report here that the NMDA-R is expressed symmetrically on the left and right right sides of the ventral hindbrain (orange arrowhead). Finally, we observed that expression of the kainate receptor was restricted to the MHB, and possibly to neurons of the posterior midbrain (Figure 2) (see next section for registration of expression patterns to specific neurons).

### Mapping of expressing neurons to the connectome

In previous studies, we have taken advantage of the small number of neurons in the *Ciona* nervous system and the stereotyped cellular anatomy to register *in situ* expression patterns to the centroids of individual neurons of the *Ciona* connectome in three-dimensions, although with varying degrees of confidence (Kourakis et al., 2019). Here, in a refinement of this approach we used complete neuron cell volumes from the connectome, rather than centroids in the registration. Briefly, 3-dimensional *in situ* image stacks were rendered into 3-dimensional expression objects with a custom Unity code, and then loaded directly into a custom Unity registration program. In the analysis, we used three VGAT/VAChT/NMDA-R, three NMDA-R/Kainate-R/VGAT, two NMDA-R/AMPA-R/VAChT, and two AMPA-R/VAChT independently-derived image stacks. Also loaded were the reconstructed neuron boundaries from the connectome. The *in situ* expression objects were first manually overlayed on the connectome cell volumes using known neurons in both datasets as anchors (Figure 3). Once the anchors were aligned, the custom Unity project was run to detect the *in situ* objects with the connectome neurons.

**Figure 3.**
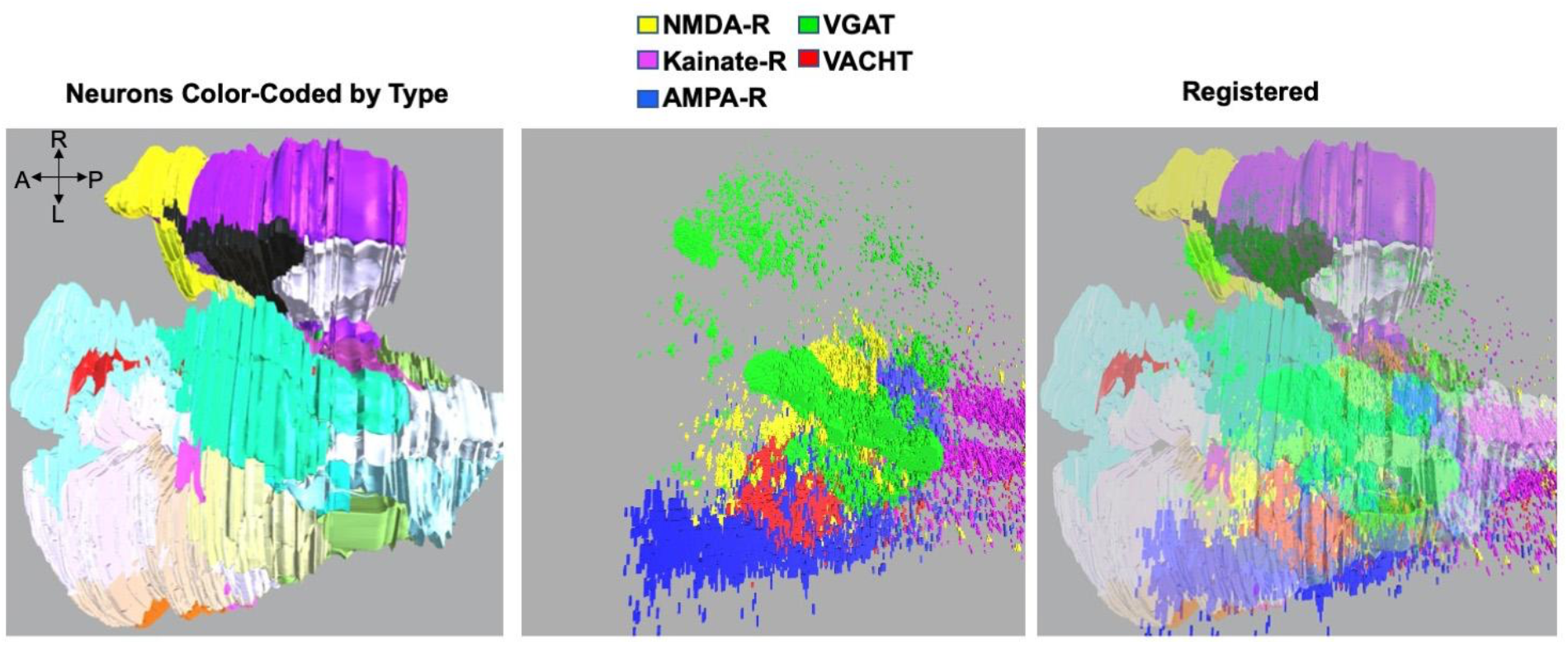
Three dimensional registration of *in situ* hybridization data to connectome neurons. The left panel shows the fore- and midbrains color-coded by type, as described (Ryan and Meinertzhagen, 2019). The center panel shows a composite of fluorescent *in situ* hybridization signals. The right panel shows the resulting registrations.

The Unity program was run for the ionotropic glutamate receptors, as well as VGAT and VAChT to generate expression predictions. The results are summarized in Table 1 and displayed as 3-dimensional overlays on the neuron centroids in Figures 4 and 5. Neurons co-expressing AMPA-R and NMDA-R were identified, as well as neurons expressing only single ionotropic receptor types, including kainate receptors. Predicted matches of glutamatergic receptor expression to neurons of the connectome in many cases conform to expectations from the connectome. For example, the expression analysis output predicts that the glutamate receptor-expressing neurons of the forebrain are *peripheral nerve interneurons* (PNINs) (Table 1), which receive direct synaptic input from VGLUT^+^ neurons - in this case from the *rostral trunk epidermal neurons* (RTENs) (Ryan et al., 2018). Thus, our prediction that glutamate-receptor-positive forebrain neurons are the neurons that receive extensive glutamatergic input supports our approach of mapping 3D *in situ* hybridization patterns to the neurons of the connectome.

**Table 1.** Predicted expression of VGAT, VAChT, AMPA-R, NMDA-R and kainate-R in neurons of the Ciona larva as given by the connectome (Ryan et al., 2016).

| Cell ID | Cell Type | VGAT+ | VAcHT | NMDAR+ | AMPA+ | KainateR+ |
| --- | --- | --- | --- | --- | --- | --- |
| 95 | aalN | 0 | + | + | 0 | 0 |
| 102 | aalN | 0 | 0 | + | 0 | 0 |
| 115 | aalN | 0 | 0 | + | 0 | 0 |
| 107 | ambiguous cells | 0 | 0 | 0 | 0 | 0 |
| 177 | ambiguous cells | 0 | 0 | 0 | 0 | 0 |
| Ant1 | Antenna | 0 | 0 | 0 | 0 | 0 |
| Ant2 | Antenna | 0 | 0 | 0 | 0 | 0 |
| 134 | AntRN | 0 | + | + | + | 0 |
| 135 | AntRN | 0 | + | + | + | 0 |
| 142 | AntRN | + | 0 | + | + | 0 |
| 143 | AntRN | 0 | 0 | + | 0 | 0 |
| 147 | AntRN | + | 0 | + | + | 0 |
| 152 | AntRN | + | 0 | + | + | + |
| 153 | AntRN | + | 0 | + | + | + |
| 159 | AntRN | + | 0 | + | + | 0 |
| 161 | AntRN | + | 0 | + | + | + |
| 120 | AntRN | 0 | 0 | + | 0 | 0 |
| 90 | Bipolar prIN | 0 | 0 | 0 | 0 | 0 |
| 92 | Bipolar prIN | 0 | 0 | 0 | 0 | 0 |
| 3 | BVIN | 0 | 0 | 0 | 0 | 0 |
| 13 | BVIN | 0 | 0 | 0 | + | 0 |
| 16 | BVIN | 0 | 0 | 0 | 0 | 0 |
| 18 | BVIN | 0 | 0 | 0 | + | 0 |
| 21 | BVIN | 0 | 0 | 0 | 0 | 0 |
| 22 | BVIN | 0 | 0 | 0 | 0 | 0 |
| 24 | BVIN | 0 | 0 | 0 | + | 0 |
| 33 | BVIN | 0 | 0 | 0 | + | 0 |
| 41 | BVIN | 0 | 0 | 0 | 0 | 0 |
| 42 | BVIN | 0 | 0 | 0 | + | 0 |
| 43 | BVIN | 0 | 0 | 0 | 0 | 0 |
| 46 | BVIN | 0 | 0 | 0 | 0 | 0 |
| 138 | BVIN | 0 | 0 | 0 | 0 | 0 |
| 1 | cor-ass BVIN | 0 | 0 | 0 | 0 | 0 |
| 2 | cor-ass BVIN | 0 | 0 | 0 | 0 | 0 |
| 15 | cor-ass BVIN | 0 | 0 | 0 | 0 | 0 |
| 17 | cor-ass BVIN | 0 | 0 | 0 | 0 | 0 |
| 23 | cor-ass BVIN | 0 | 0 | 0 | 0 | 0 |
| 38 | cor-ass BVIN | 0 | 0 | 0 | 0 | 0 |
| 48 | cor-ass BVIN | 0 | 0 | 0 | + | 0 |
| 50 | cor-ass BVIN | 0 | 0 | 0 | 0 | 0 |
| 55 | cor-ass BVIN | 0 | 0 | 0 | + | 0 |
| 59 | cor-ass BVIN | 0 | 0 | 0 | + | 0 |
| 60 | cor-ass BVIN | 0 | 0 | 0 | + | 0 |
| 62 | cor-ass BVIN | 0 | 0 | 0 | 0 | 0 |
| 68 | cor-ass BVIN | 0 | 0 | + | 0 | 0 |
| 70 | cor-ass BVIN | 0 | 0 | 0 | 0 | 0 |
| 73 | cor-ass BVIN | 0 | 0 | 0 | + | 0 |
| 78 | cor-ass BVIN | 0 | 0 | 0 | 0 | 0 |
| 79 | cor-ass BVIN | 0 | 0 | + | 0 | 0 |
| coronet1 | Coronet | 0 | 0 | 0 | 0 | 0 |
| coronet 10 | Coronet | 0 | 0 | + | + | 0 |
| coronet11 | Coronet | 0 | 0 | + | + | 0 |
| coronet12 | Coronet | 0 | 0 | 0 | 0 | 0 |
| coronet13 | Coronet | 0 | 0 | 0 | 0 | 0 |
| coronet14 | Coronet | 0 | 0 | 0 | 0 | 0 |
| coronet15 | Coronet | 0 | 0 | 0 | 0 | 0 |
| coronet16 | Coronet | 0 | 0 | 0 | 0 | 0 |
| coronet2 | Coronet | 0 | 0 | 0 | + | 0 |
| coronet3 | Coronet | 0 | 0 | 0 | 0 | 0 |
| coronet4 | Coronet | 0 | 0 | 0 | + | 0 |
| coronet5 | Coronet | 0 | 0 | 0 | 0 | 0 |
| coronet6 | Coronet | 0 | 0 | 0 | + | 0 |
| coronet7 | Coronet | 0 | 0 | 0 | 0 | 0 |
| coronet8 | Coronet | 0 | 0 | 0 | 0 | 0 |
| coronet9 | Coronet | 0 | 0 | + | + | 0 |
| 109 | Eminens | + | 0 | 0 | 0 | 0 |
| 99 | Eminens | + | 0 | 0 | 0 | 0 |
| 103 | non-sensory RN | + | 0 | 0 | 0 | 0 |
| 106 | non-sensory RN | 0 | 0 | 0 | + | 0 |
| 122 | non-sensory RN | + | 0 | 0 | 0 | 0 |
| 125 | non-sensory RN | 0 | + | + | + | 0 |
| 93 | non-sensory RN | 0 | 0 | 0 | + | 0 |
| 160 | PBV PNIN | + | 0 | 0 | 0 | + |
| 162 | PBV PNIN | + | 0 | 0 | + | + |
| 163 | PBV PNIN | + | 0 | 0 | 0 | + |
| 164 | PBV PNIN | 0 | 0 | 0 | 0 | + |
| 20 | PNIN | 0 | 0 | 0 | 0 | 0 |
| 25 | PNIN | 0 | 0 | 0 | 0 | 0 |
| 29 | PNIN | 0 | 0 | 0 | 0 | 0 |
| 30 | PNIN | 0 | 0 | 0 | 0 | 0 |
| 4 | PNIN | 0 | 0 | 0 | 0 | 0 |
| 6 | PNIN | 0 | 0 | 0 | 0 | 0 |
| 85 | PNIN | + | 0 | 0 | 0 | 0 |
| 61 | PNIN | + | 0 | 0 | 0 | 0 |
| 65 | PNIN | + | 0 | 0 | 0 | 0 |
| 88 | PNIN | + | 0 | 0 | 0 | 0 |
| 131 | PN RN | + | 0 | 0 | 0 | 0 |
| prIII-101 | PR (III) | 0 | 0 | 0 | 0 | 0 |
| prIII-110 | PR (III) | 0 | 0 | 0 | 0 | 0 |
| prIII-113 | PR (III) | 0 | 0 | 0 | 0 | 0 |
| prIII-114 | PR (III) | 0 | 0 | 0 | 0 | 0 |
| prIII-6 | PR (III) | 0 | 0 | 0 | 0 | 0 |
| prIII-7 | PR (III) | 0 | 0 | 0 | 0 | 0 |
| prIII-84 | PR (III) | 0 | 0 | 0 | 0 | 0 |
| 108 | pr-AMG RN | 0 | 0 | 0 | 0 | 0 |
| 116 | pr-AMG RN | + | 0 | + | 0 | 0 |
| 124 | pr-AMG RN | + | 0 | 0 | 0 | 0 |
| 127 | pr-AMG RN | 0 | 0 | + | 0 | 0 |
| 140 | pr-AMG RN | 0 | + | + | + | 0 |
| 157 | pr-AMG RN | + | 0 | + | + | 0 |
| 74 | pr-AMG RN | 0 | 0 | 0 | 0 | 0 |
| 94 | pr-AMG RN | + | 0 | + | 0 | 0 |
| 123 | pr-BTN RN | + | 0 | + | 0 | 0 |
| 130 | pr-BTN RN | + | 0 | + | 0 | 0 |
| 105 | pr-cor RN | 0 | + | + | + | 0 |
| 112 | pr-cor RN | 0 | + | + | + | 0 |
| 119 | pr-cor RN | 0 | + | + | + | 0 |
| 100 | prRN | 0 | + | + | + | 0 |
| 121 | prRN | 0 | + | + | + | 0 |
| 126 | prRN | 0 | + | + | + | 0 |
| 80 | prRN | 0 | 0 | 0 | 0 | 0 |
| 86 | prRN | 0 | + | + | + | 0 |
| 96 | prRN | 0 | + | + | + | 0 |
| AMG1 | AMG | + | 0 | 0 | 0 | 0 |
| AMG2 | AMG | + | 0 | 0 | 0 | 0 |
| AMG3 | AMG | + | 0 | 0 | 0 | 0 |
| AMG4 | AMG | + | 0 | 0 | 0 | 0 |
| AMG5 | AMG | 0 | + | 0 | 0 | 0 |
| AMG6 | AMG | 0 | 0 | 0 | 0 | 0 |
| AMG7 | AMG | 0 | 0 | 0 | 0 | 0 |
| ddNL | ddN | 0 | + | + | + | 0 |
| ddNR | ddN | 0 | + | + | 0 | 0 |
| MGIN1L | MGIN | 0 | + | + | + | 0 |
| MGIN1R | MGIN | 0 | + | + | 0 | 0 |
| MGIN2L | MGIN | 0 | + | + | + | 0 |
| MGIN2R | MGIN | 0 | + | + | 0 | 0 |
| MGIN3L | MGIN | 0 | + | + | + | 0 |
| MGIN3R | MGIN | 0 | + | + | 0 | 0 |
| MN1L | MN | 0 | + | + | + | 0 |
| MN1R | MN | 0 | + | + | 0 | 0 |
| MN2L | MN | 0 | + | 0 | + | 0 |
| MN2R | MN | 0 | + | 0 | 0 | 0 |
| MN3L | MN | 0 | + | + | + | 0 |
| MN3R | MN | 0 | + | + | 0 | 0 |
| 165 | Neck | 0 | + | + | + | + |
| 166 | Neck | 0 | + | + | 0 | + |
| pr1 | PR (I) | 0 | 0 | 0 | 0 | 0 |
| pr10 | PR (I) | + | 0 | 0 | 0 | 0 |
| pr11 | PR (I) | 0 | 0 | 0 | 0 | 0 |
| pr12 | PR (I) | 0 | 0 | 0 | 0 | 0 |
| pr13 | PR (I) | 0 | 0 | 0 | 0 | 0 |
| pr14 | PR (I) | 0 | 0 | 0 | 0 | 0 |
| pr15 | PR (I) | 0 | 0 | 0 | 0 | 0 |
| pr16 | PR (I) | 0 | 0 | 0 | 0 | 0 |
| pr17 | PR (I) | 0 | 0 | 0 | 0 | 0 |
| pr18 | PR (I) | 0 | 0 | 0 | 0 | 0 |
| pr19 | PR (I) | 0 | 0 | 0 | 0 | 0 |
| pr2 | PR (I) | 0 | 0 | 0 | 0 | 0 |
| pr20 | PR (I) | 0 | 0 | 0 | 0 | 0 |
| pr21 | PR (I) | 0 | 0 | 0 | 0 | 0 |
| pr22 | PR (I) | 0 | 0 | 0 | 0 | 0 |
| pr23 | PR (I) | 0 | 0 | 0 | 0 | 0 |
| pr3 | PR (I) | 0 | 0 | 0 | 0 | 0 |
| pr4 | PR (I) | 0 | 0 | 0 | 0 | 0 |
| pr5 | PR (I) | 0 | 0 | 0 | 0 | 0 |
| pr6 | PR (I) | 0 | 0 | 0 | 0 | 0 |
| pr7 | PR (I) | 0 | 0 | 0 | 0 | 0 |
| pr8 | PR (I) | 0 | 0 | 0 | 0 | 0 |
| pr9 | PR (I) | 0 | 0 | 0 | 0 | 0 |
| pr-a | PR (II) | + | 0 | 0 | 0 | 0 |
| pr-b | PR (II) | + | 0 | 0 | 0 | 0 |
| pr-c | PR (II) | + | 0 | 0 | 0 | 0 |
| pr-d | PR (II) | + | 0 | 0 | 0 | 0 |
| pr-e | PR (II) | + | 0 | 0 | 0 | 0 |
| pr-f | PR (II) | + | 0 | 0 | 0 | 0 |
| pr-g | PR (II) | + | 0 | 0 | 0 | 0 |

**Figure 4.**
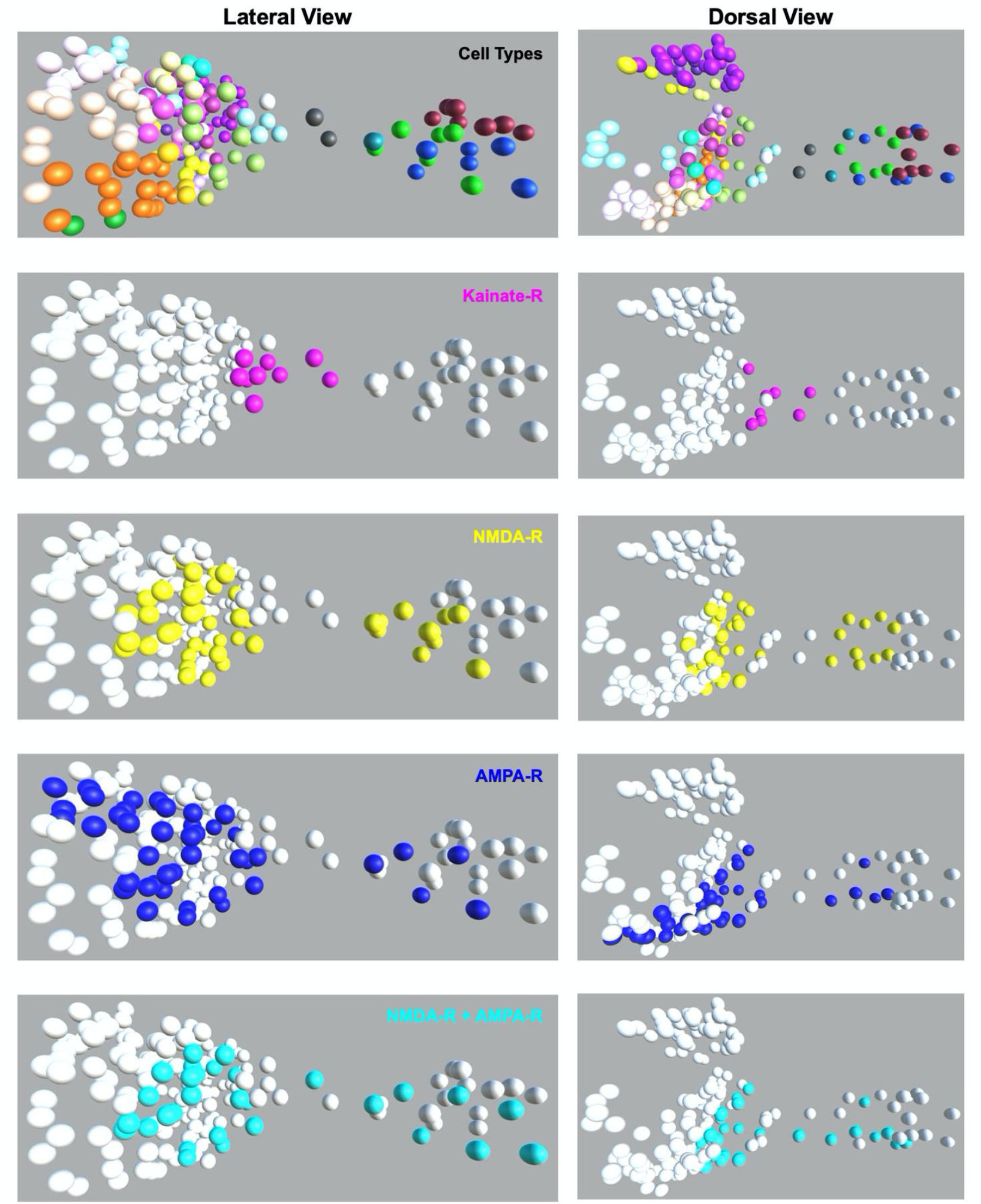
Summary of glutamatergic receptor expression. The two top panels show forebrain to hindbrain neuron centroids given by (Ryan et al., 2016), and are colored by neuron class according to (Ryan and Meinertzhagen, 2019), see SFigure 1. The bottom panels show the predicted distribution of AMPA, NMDA and kainate receptors at larval stage.

**Figure 5.**
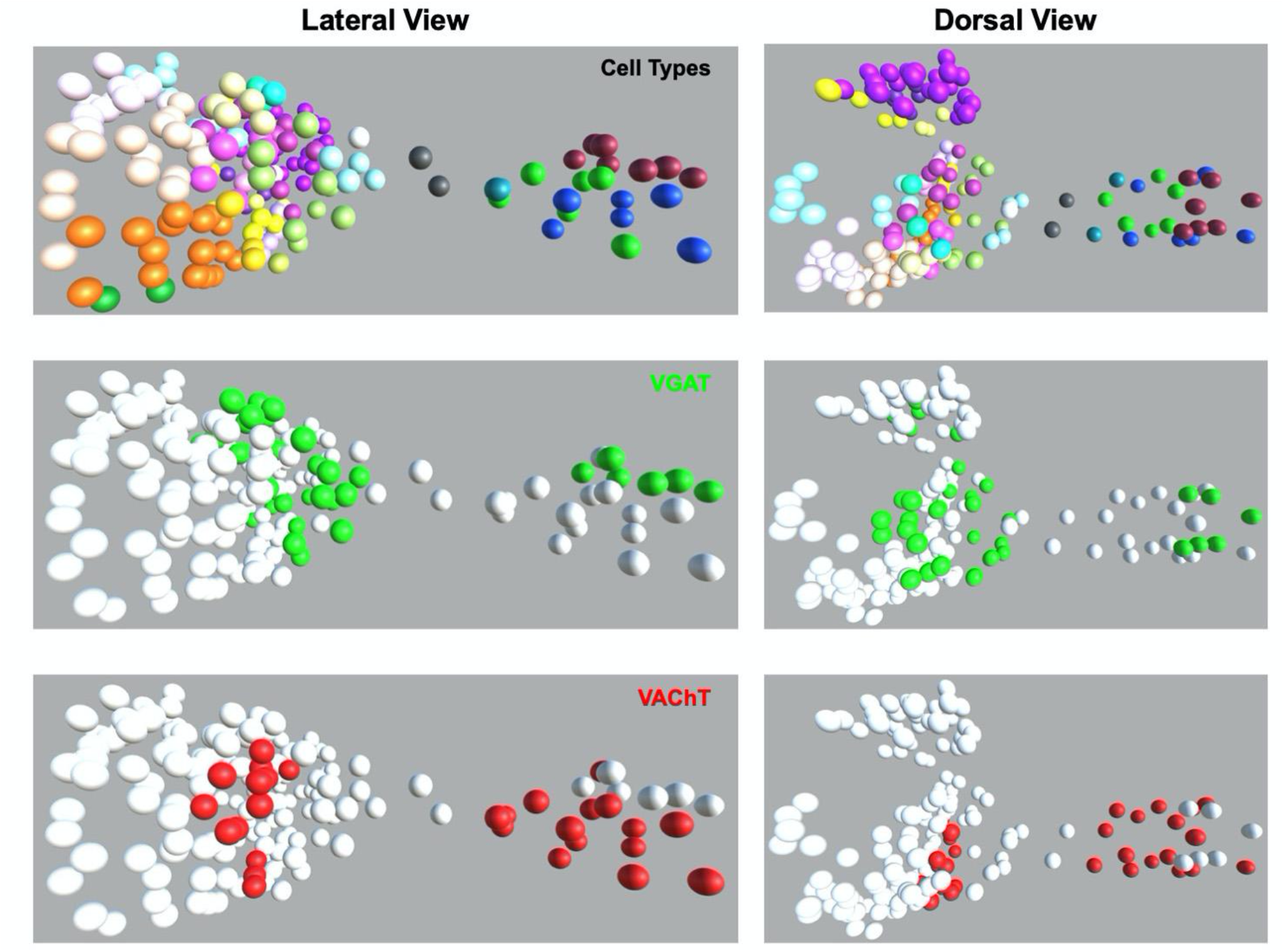
Summary of VAChT and VGAT expression. The two top panels show forebrain to hindbrain neuron centroids given by (Ryan et al., 2016), and are colored by neurons class according to (Ryan and Meinertzhagen, 2019), see SFigure 1. The bottom panels show the predicted distribution of VAChT and VGAT at the larval stage.

### Expression of mGlu receptors in the peripheral nervous system

Analysis of the *Ciona* genome revealed the presence of three putative metabotropic glutamate receptors (mGluR) (Kamesh et al., 2008). Based on their orthologies to vertebrate mGluRs, they were named mGluR123, mGluR478, and mGluR147. Our analysis of a previously published single-cell RNAseq (scRNAseq) dataset (Cao et al., 2019) indicated that mGluR123 was expressed at higher levels at the larval stage than the other two mGluRs (SFigure 2). *In situ* hybridization for mGluR123 revealed expression in the epidermal sensory neurons (ESNs), but no apparent central nervous system expression (Figure 6). This observation agrees with distribution of mGluR123 transcripts among the tissue types for the scRNAseq dataset (Figure S3). While mGluR123 expression was widespread among the ESNs, including the rostral trunk epidermal neurons (RTEN), the posterior rostral trunk epidermal neurons (pATEN), and the dorsal and ventral caudal epidermal neurons (DCEN and VCEN, respectively), we did not observe expression in the anterior rostral trunk epidermal neurons, which are found between the RTENs and the pATENs [Figure 6; for a complete description of *Ciona* epidermal sensory neurons refer to (Ryan et al., 2018)]. While the 3D coordinates for the ENS were not included in the connectome dataset, the ESNs are well described and easily identified, allowing *in situ* expression patterns to be confidently attributed. An *in situ* hybridization was also performed for mGluR478, and no expression was detected - consistent with the low expression indicated by the scRNAseq dataset (SFigure 2). The scRNAseq dataset indicates that mGluR147 is also expressed at a lower level than mGluR123, and no *in situ* hybridization was attempted. However, the tissue distribution of mGluR147 is similar to that of mGluR478, with expression concentrated in the epidermis (SFigure 2).

**Figure 6.**
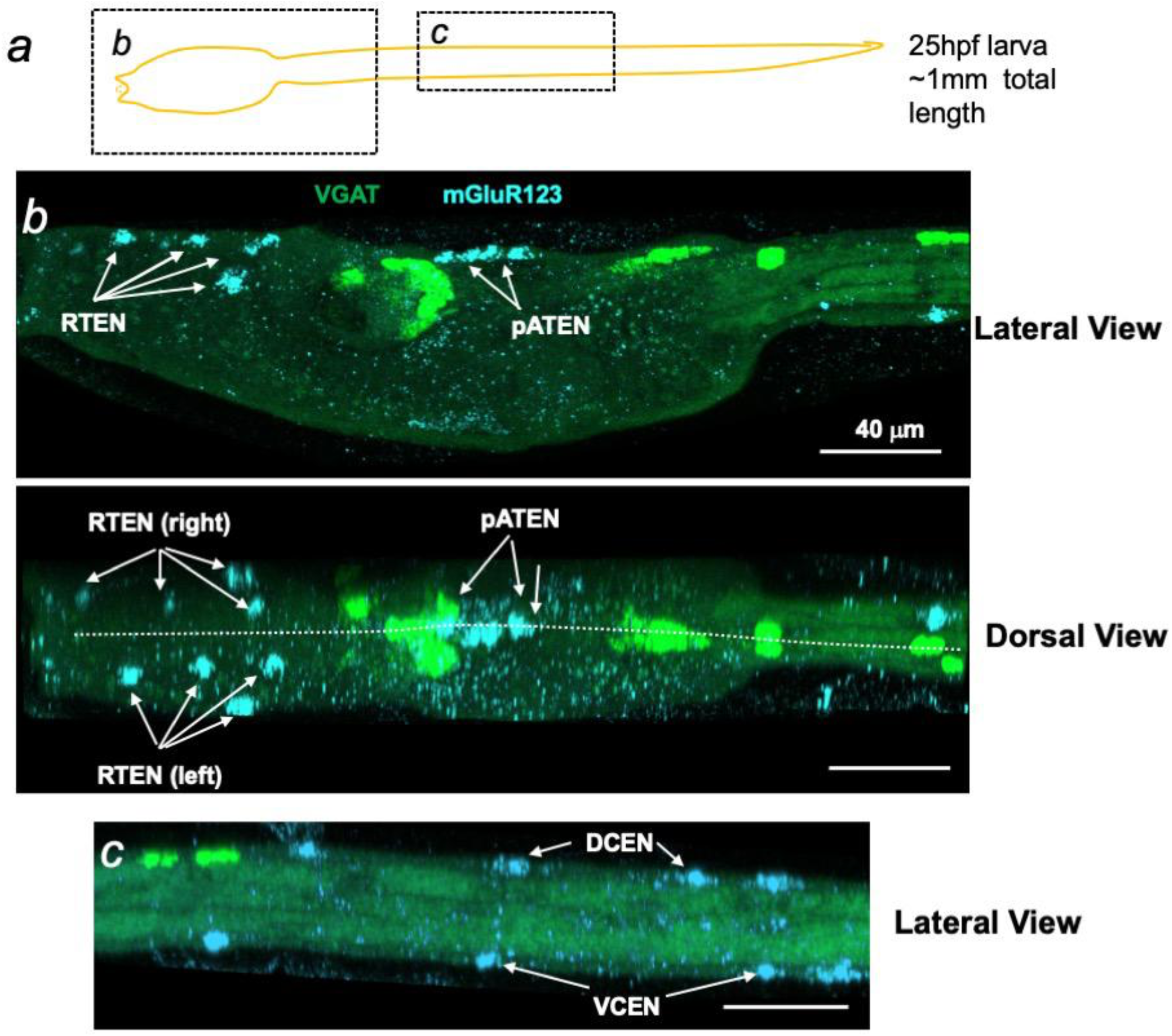
Expression of mGluR123 by *in situ* hybridization. **a.** Cartoon of *Ciona* larva indicating regions shown in panels b and c. **b.** mGluR123 expression in the trunk of a *Ciona* larva. Arrows point to mGluR123-expressing epidermal sensory neurons. Abbreviations: RTEN, rostral trunk epidermal neurons; aATEN, anterior apical trunk epidermal neurons; pATEN posterior apical trunk epidermal neurons. Both dorsal and lateral views are shown. The dotted line in the dorsal view indicates the midline. **c**. mGluR123 expression in the tail of a *Ciona* larva. Abbreviations: DCEN, dorsal caudal epidermal neurons; VCEN and ventral caudal epidermal neurons.

### NMDA receptors are required for sensorimotor responses

In a previous study we found that the AMPA-R inhibitor perampanel blocked negative phototaxis, the active swimming of *Ciona* larvae away from a light source, but not their ability to respond to dimming light (Kourakis et al., 2019). This result is consistent with the differential expression of AMPA-R on the primary interneuron targets of the photoreceptors. The prRNs are thought to mediate phototaxis, and express both AMPA-R and NMDA-R (Table 1). By contrast, the pr-AMG RNs are thought to mediate the dimming response, and a subset is predicted to express NMDA-R but not AMPA-R (Table 1).

The non-competitive NMDA receptor antagonist MK801 was used to assess the role of NMDA-Rs in visuomotor responses. MK801 shows strong inhibition and specificity of NMDA-Rs in both vertebrates and invertebrates (Vogeler et al., 2021; Wong et al., 1986). The effect of NMDA-R inhibition was first tested in a dimming assay (Salas et al., 2018). Larvae were recorded using far-red illumination (700 nm), while a 505 nm LED lamp was dimmed from 3 to 0.3 mW/cm^2^ midway through the recording. We observed that unlike the AMPA-R antagonist perampanel, MK801 completely inhibited the dimming response (SMovie 1). Figure 7 shows temporal projections of the movies for the 5 seconds immediately before and after the dim. Before the dim, larvae are mostly stationary, but in response to dimming the control larvae immediately initiate swimming, which are seen as lines in the time-projection image (see labels in Figure 7). By contrast, larvae treated with 0.5 mM MK801 did not respond to dimming (bottom two panels). A similar result was observed in a phototaxis assay (SMovie 2). In the phototaxis assay, larvae are placed in a petri dish with a light source of constant intensity from one side and recorded for one hour (Salas et al., 2018). At the end of the assay period, phototaxis is evident by the accumulation of larvae at the side of the petri dish furthest from the light source. Figure 8 shows images from SMovie 2 of the control and MK108-treated larvae at the start of the phototaxis assay (t=0), in which the larvae can be seen evenly distributed across the petri dishes (left panels). The right panels show a temporal projection of the movie from time points 30 to 60 minutes. The accumulation of larvae at the left side of the petri dish (away from the light) is evident in the control, but not in the MK801-treated sample. Thus the NMDA-R antagonist MK801 is effective at inhibiting both the dimming response and phototaxis, unlike the AMPA antagonist perampanel, which only inhibits phototaxis. Because of the widespread expression of NMDA-Rs, including the mid- and hindbrain (Table 1 and Figure 4), it is not possible to attribute the behavioral effects of MK801 to particular neurons of the circuit.

**Figure 7.**
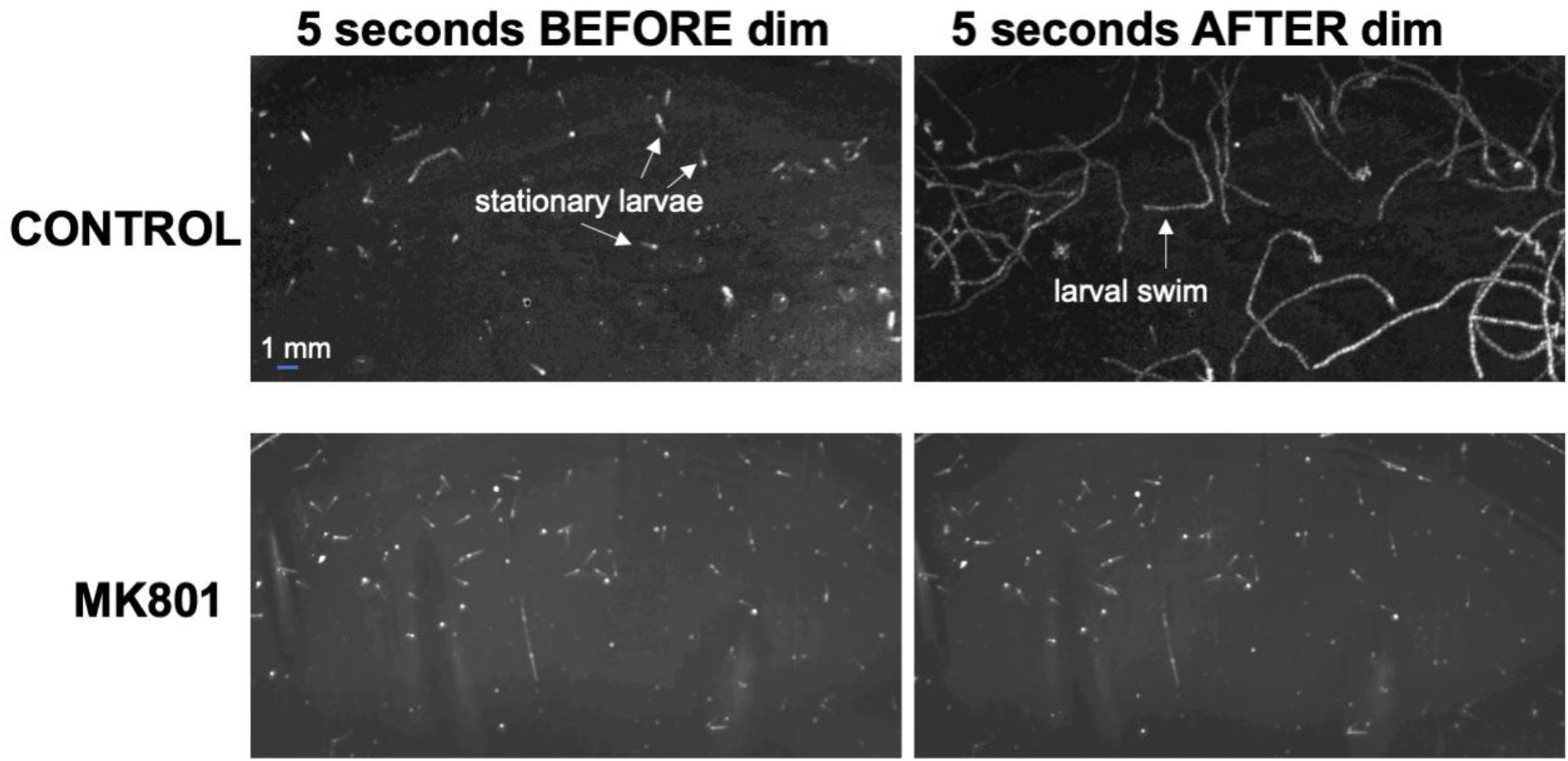
MK801 blocks the dimming response. The left two panels show *Ciona* larvae in a temporal projection of the 5 seconds preceding light dimming for both control and MK801-treated larvae. Lines represent larval paths within the 5 second projection. Most larvae are stationary in the five seconds preceding the dimming event. In the five seconds following light dimming the CONTROL larvae are observed swimming (white lines), while the MK801-treated larvae do not respond.

**Figure 8.**
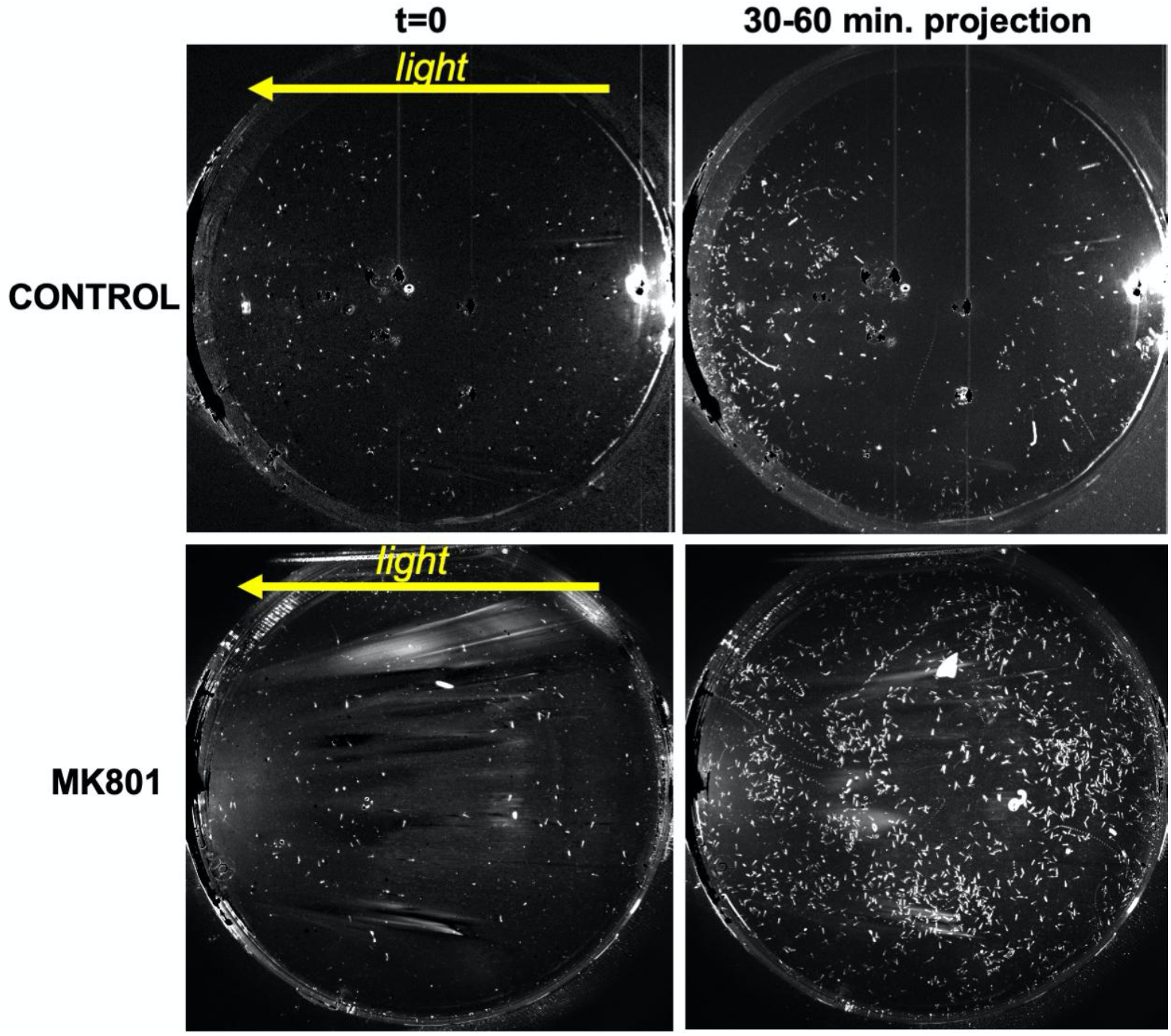
MK801 blocks the phototaxis response. The left two panels show the first frame of the phototaxis assay with the yellow arrow indicating the direction of the light. The right two panels show temporal projects from 30 minutes to 60 minutes in directional light. Larvae can be seen accumulated on the left in the control and more dispersed in the MK801-treated.

## Discussion

The small number of neurons in the *Ciona* larva, together with the published connectome and easily quantified sensorimotor behaviors, make it a powerful model for neuron circuit analysis. Moreover, *Ciona* is currently the only chordate for which a complete analysis of all neurons in a nervous system is possible. Key to making testable models of neural circuits is a knowledge of the distribution of neurotransmitter receptors. We present here a comprehensive analysis at the individual neuron level of glutamate receptor expression in the *Ciona* larval nervous system. Also presented are analyses for the distribution of VGAT^+^ and VAChT^+^ neurons, which expands our previous report (Kourakis et al., 2019) to additional neurons in the CNS, and resolves the neurotransmitter use of neurons that were previously ambiguous. The challenge of mapping gene expression data to the connectome is that the fully annotated connectome exists for only a single individual, while *in situ* hybridization results can be generated from multiple individuals. Our ability to map expression patterns to individual neurons of the connectome with confidence would only be possible if the CNS showed a high degree of stereotypy. The issue of stereotypy is addressed in a previous publication (Kourakis et al., 2019), in which we show that the larval CNS is highly stereotyped, with a few notable exceptions - most prominently the a*scending contralateral inhibitory neurons* of the hindbrain. It is also important to note that our results are for a specific developmental stage of the larva (∼25 hours post-fertilization at 18°C). We have observed changes in *Ciona* larval behavior over time-scales as short as 2-3 hours (Bostwick et al., 2020; Salas et al., 2018), so it is possible that the expression of the glutamate receptors is temporally dynamic within the larva. Nevertheless, in the temporal window that was analyzed here, the distribution of glutamate receptors in the *Ciona* nervous system provides new insight into neural circuits driving sensorimotor behaviors.

### Glutamate receptors in the fore- and midbrain

We previously reported using VGLUT as a marker for glutamatergic neurons, and observed expression only in sensory neurons (photoreceptors, antenna cells and peripheral sensory neurons), with no apparent glutamatergic interneurons (Kourakis et al., 2019). The expression of AMPA-R and NMDA-R reported here agrees with the synaptic targeting of fore- and midbrain neurons by VGLUT^+^ sensory neurons. Interestingly, while most photoreceptors are glutamatergic, and the connectome predicts extensive chemical synapses between them, we found no evidence of glutamate receptor expression in the photoreceptors. The nature of the apparent chemical synapses between the photoreceptors remains to be determined.

In contrast to the forebrain, which only expresses glutamate receptors in a limited number of neurons (specifically the PNINs; Table 1), expression of AMPA-R and NMDA-R in the midbrain was much more extensive. The anterior midbrain receives input from the photoreceptors, while the posterior midbrain receives synaptic input from the otolith-associated *antennae sensory neurons*, which mediate gravitaxis (Bostwick et al., 2020; Ryan et al., 2016). The target of the two VGLUT^+^ antenna neurons - the VGAT^+^ *antenna relay neurons* (AntRNs), all express either NMDA-R, AMPA-R, or both (Table 1; Figures 4 and 5). The eleven AntRNs have a surprisingly complex synaptic connectivity, with some receiving input from one antenna neuron, some from the other, and some from both (Ryan et al., 2016). We previously reported that perampanel was effective at blocking gravitaxis at 21 hpf, but not at 25 hpf, leading us to speculate, based on the presence of extensive gap junctions between the antenna cells and AntRNs, the synapse matured from chemical to electrical during that temporal window (Bostwick et al., 2020). The results here which suggest heterogeneity of glutamate receptor expression only add to the apparent complexity, and call for further investigation.

Photoreceptor input to the *Ciona* larval midbrain is even more complex than antenna cell input. The visual organ of *Ciona*, the ocellus, contains two distinct groups of photoreceptors, PR-I and PR-II (Horie et al., 2008b; Kourakis et al., 2019). PR-I consists of 23 photoreceptors (21 VGLUT^+^, one VGAT^+^, and one VGLUT^+^/VGAT^+^), all of which project their outer segments into the ocellus pigment cell. The direction-dependent shading of the photoreceptors by the pigment cell as larvae perform short orienting swims provides a cue to the direction of light, and thereby mediates negative phototaxis (Salas et al., 2018). By contrast, the seven PR-II photoreceptors (three VGAT^+^ and four VGLUT^+^/VGAT^+^) are not associated with the pigment cell and are sensitive to light from all directions, and thereby mediate a light-dimming behavior (Salas et al., 2018). The PR-I photoreceptors project axons to two distinct classes of primary interneurons in the midbrain, the prRNs and the pr-AMG RNs (Figure 9a; Ryan et al., 2016). At least five of the six prRNs are predicted to be cholinergic and express both AMPA-R and NMDA-R (Table1; Figure 4). Our model of phototaxis hypothesizes that the cholinergic prRNs send excitatory input from the VGLUT^+^ photoreceptors to secondary cholinergic interneurons in the hindbrain, and then to the motor neurons (Kourakis et al., 2019). However, the other midbrain interneuron class targeted by the PR-I photoreceptors, the pr-AMG RNs are predicted to be mostly inhibitory (*i.e*., VGAT^+^; Table 1 and Figure 5). The prediction that PR-I photoreceptors project to both excitatory (prRNs) and inhibitory (pr-AMG RNs) relay neurons initially appears to be difficult to account for within the model. However, our observation here that the pr-AMG RNs express NMDA-Rs, but not AMPA-Rs, provides an explanation (Figure 9). Namely, AMPA-Rs mediate fast excitatory responses while NMDA-Rs have a slower modulatory role. Moreover, we hypothesized previously that the projection of the PR-I photoreceptors to both excitatory and inhibitory primary interneurons constitute an *incoherent feedforward loop* circuit motif that functions in visual processing to generate the observed fold-change detection behavior in phototaxis (Borba et al., 2021). In fold-change detection the response scales with the magnitude of the temporal change in sensory input, not with the absolute value of the input (Adler and Alon, 2018). The presence of NMDA-Rs, but not AMPA-Rs, on the pr-AMG RNs fits well with a hypothesized modulatory role in this circuit.

**Figure 9.**
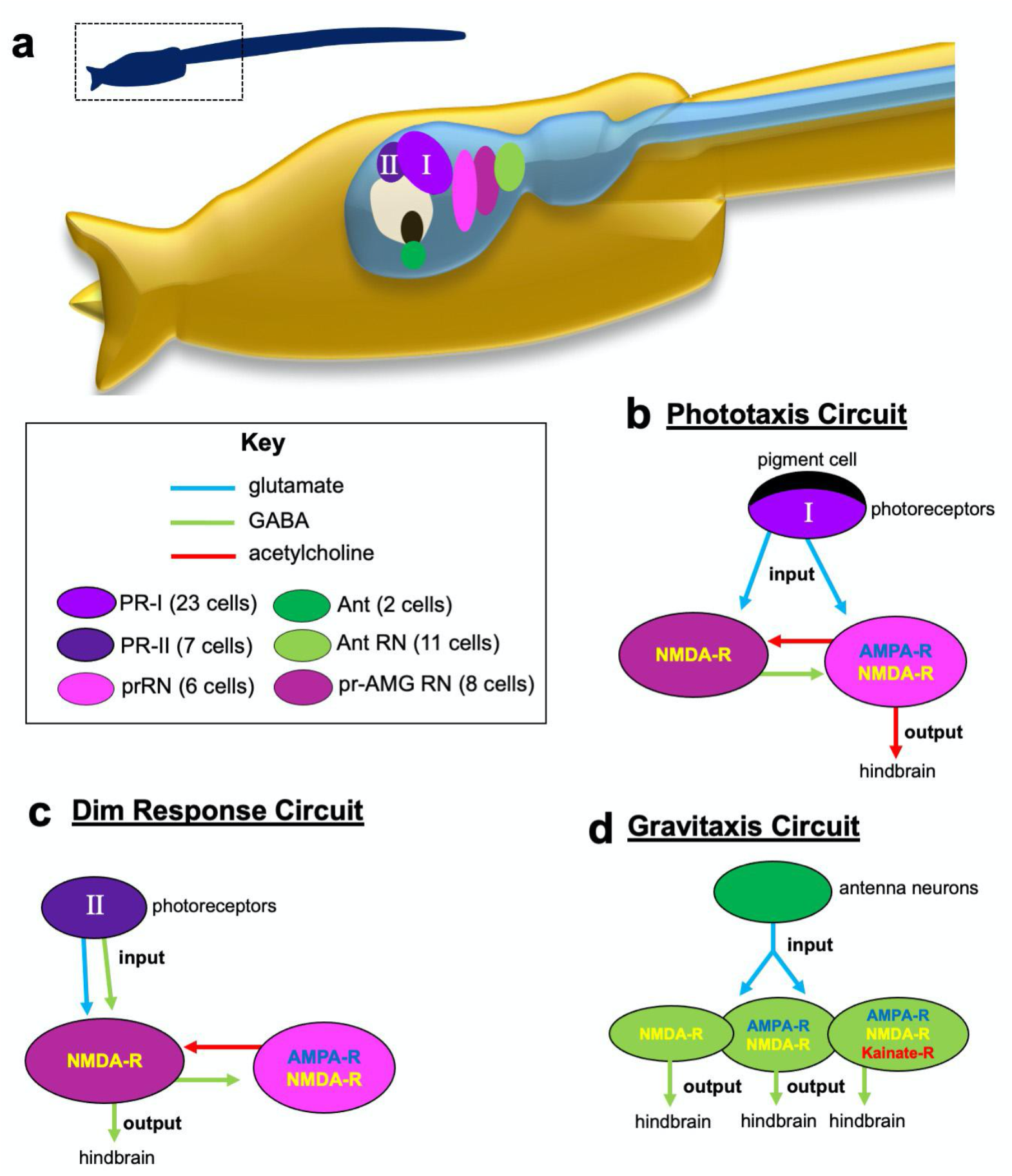
Model neural circuits with contributions by glutamate receptors. **a**. Overview of central nervous system visual and gravitactic circuits. The left and right centers in the hindbrain contain secondary interneurons and motor neurons (not shown). **b**. Phototaxis circuit. Note that the PR-I photoreceptors (input) project to both the prRNs and the pr-AMG RNs, but that output is exclusively from the prRNs. **c**. Dim response circuit. This response involves the PR-II photoreceptors, which project exclusively to the pr-AMG RNs. **d**. Gravitaxis circuit. The gravity-sensitive antenna neurons project to VGAT^+^ primary interneurons that are heterogeneous in their expression of ionotropic glutamate receptors. In all panels, neuron classes are named and color-coded according to (Ryan and Meinertzhagen, 2019). Abbreviations: OT= otolith; PR-I = photoreceptor group I; PR-II = photoreceptor group II; prRN = photoreceptor relay neurons; pr-AMG RN= photoreceptor-ascending MG neuron relay neurons Ant= antenna neurons; Ant RN = antenna relay neuron.

The PR-II photoreceptors project only to the pr-AMG RNs (Figure 9b). All of the PR-II photoreceptors are VGAT^+^ (and thus likely inhibitory), with a subset being dual VGLUT^+^/VGAT^+^. Because their sole synaptic targets, the pr-AMG RNs, also appear to be mostly inhibitory, we hypothesized that the dimming response is mediated by disinhibition, a hypothesis supported by pharmacology (Kourakis et al., 2019). However, the dual release of glutamate and GABA by a subset of the PR-II photoreceptors was not easily accounted for in the disinhibition model. Our finding here that a set of the pr-AMG RNs express NMDA-R, but not AMPA-Rs (Table 1, Figure 4), agrees well with the disinhibition model. In other words, glutamate could serve a modulatory role in the dimming response circuit, perhaps acting via GABA receptors, as has been shown previously (Marsden et al., 2007). Moreover, like the phototaxis behavior, the dimming response shows fold-change detection, although with a different putative circuit motif than in the phototaxis circuit (Borba et al., 2021), and modulation of GABA receptors could serve as the “memory” component of the fold-change detection circuit, as has been described for other systems showing fold-change detection (Lyashenko et al., 2020).

### Glutamate receptors in the hindbrain

We previously reported that AMPA-R transcripts are present asymmetrically in the hindbrain, with expression only observed on the left side (Kourakis et al., 2021). By contrast, we report here that NMDA-R transcripts were observed equally in the left and right hindbrain. The presence of glutamate receptors in the hindbrain initially appears to be paradoxical, as none of the AMPA-R and NMDA-R expressing hindbrain neurons receive direct sensory input (Ryan et al., 2018, 2016), and thus no direct glutamatergic synaptic input. By contrast, the AMG neurons of the dorsal hindbrain are primary synaptic targets of the glutamatergic peripheral sensory neurons (Ryan et al., 2018), yet have no detectable glutamate receptor expression (Table 1). However, the connectome predicts extensive electrical synapses between the peripheral sensory neurons and the AMGs, suggesting the transmission to the AMGs is not chemical. Nevertheless, since the peripheral sensory neurons are the only glutamatergic neurons to enter the hindbrain, we speculate that they may signal to the glutamate receptor-expressing hindbrain neurons extrasynaptically, which has been observed for glutamate signaling, and the distance between the peripheral sensory neuron termini and their putative hindbrain targets is well within the diffusion range of glutamate (Pál, 2018). *Kainate-R and mGluR expressions.* In addition to investigating the expression of AMPA-R and NMDA-R, we also examined kainate and metabotropic glutamate receptors. Kainate receptors are ionotropic and appear to have functions both pre- and post-synaptically (Contractor et al., 2011). We observed kainate-R expression in a distinct set of neurons in the posterior midbrain and neck (Table 1 and Figure 4). While we predict that three of the AntRNs express kainate-R, the function of the other kainate-R^+^ neurons is not known. Of the three predicted metabotropic glutamate receptors (mGlu-R), we only found evidence for expression of one of them (SFigure 2), and only in peripheral sensory neurons (Figure 8). The function of this mGlu-R is not known, but we speculate that it may play a role in signaling between the ESNs, and possibly mediate attenuation of the touch response.

### Behavioral requirements for NMDA-R and AMPA-R

Our observation that treatment of larvae with MK801 blocked visuomotor responses (phototaxis and dim response) came as a surprise. Because of the documented modulatory role of NMDA-Rs, we speculate that tonic activation of NMDA-R, perhaps extra synaptically in some cases, may be necessary to maintain sensorimotor responses. Because of the widespread distribution of NMDA-Rs in the CNS, including in motor neurons and MGINs, which are common to all sensorimotor circuits, it is not possible to attribute the result to blocking NMDA-Rs in a particular neuron class. Neuron-specific targeting methods, such as CRISPR, will be required to assess the role of NMDA-Rs in particular circuits.

## Acknowledgments

We thank Kerrianne Ryan for helpful discussion and providing the connectome cell meshes. We also thank the team members of ANISEED for providing tools to help in analysis of glutamate receptors.

## Data Availability

*In situ* hybridization datasets are available at https://www.brainimagelibrary.org/. Expression data mapped to the connectome is available at https://morphonet.org/2Tn9x6ZF.

