## Supplemental Files for "A whole nervous system atlas of glutamate receptors reveals distinct receptor roles in sensorimotor circuits"

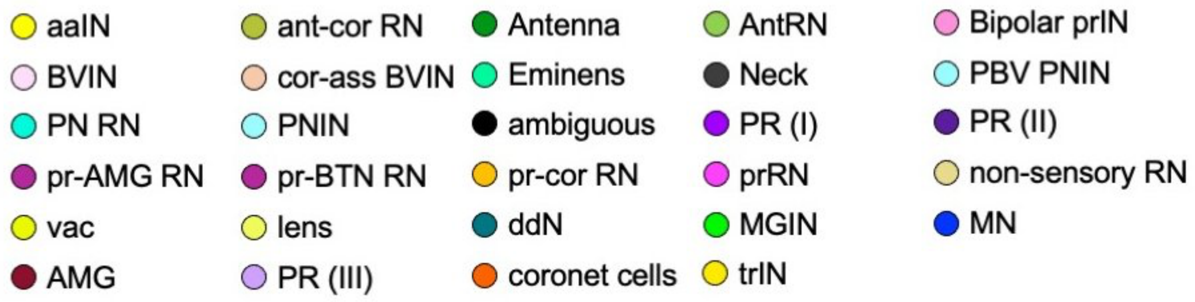


**SFigure 1**. Key for color-coded neuron types in Figures 4 and 5.

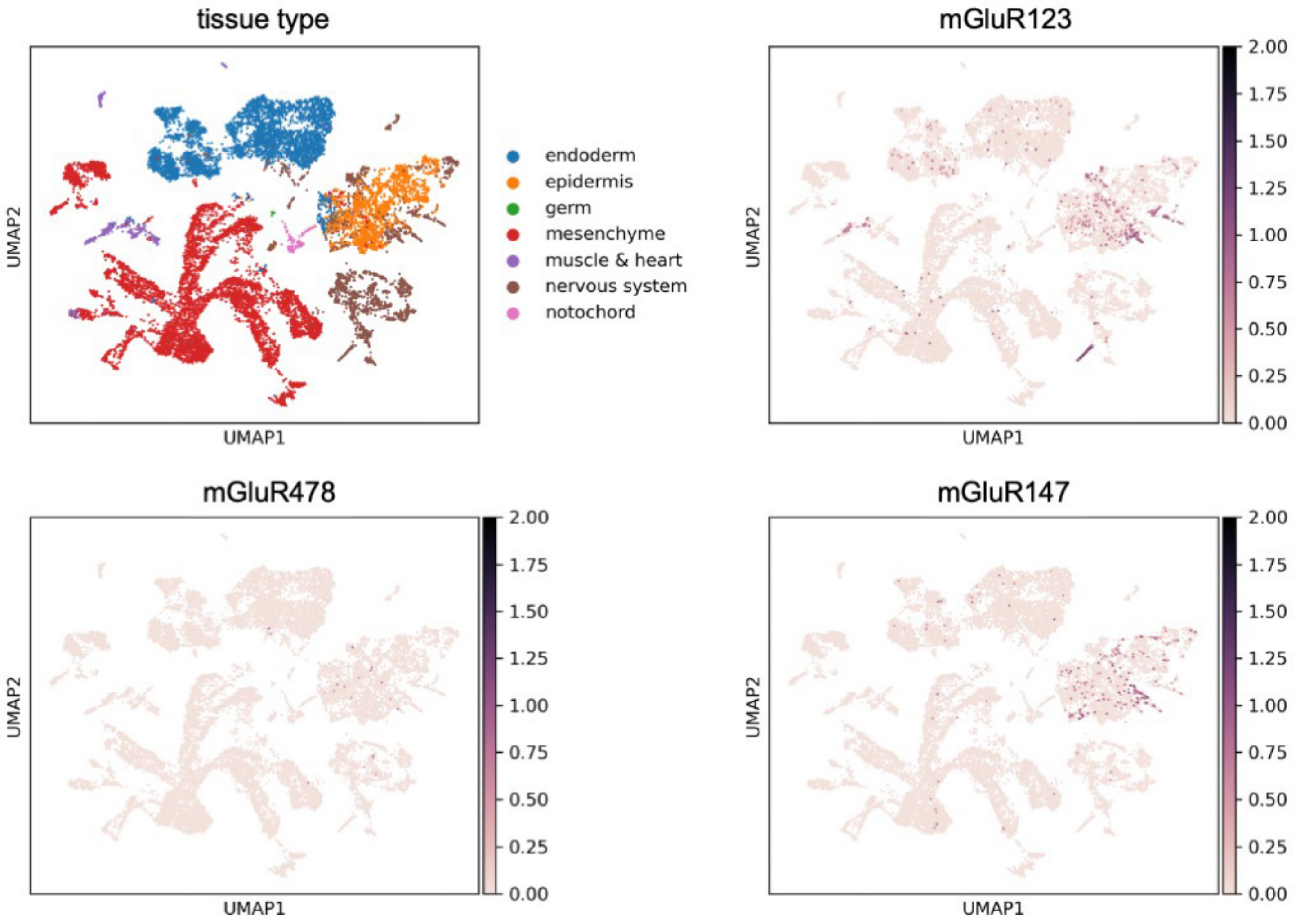


**SFigure 2.** Expression of mGlu receptors (mGluR) in the *Ciona* larvae single cell RNaseq dataset from (Cao et al., 2019). The top left panel shows cells clustered by UMAP analysis and color-coded by tissue type. The remaining three panels show the distribution of the three putative mGlu receptors in the clusters. Notice that mGluR123 is more highly expressed than the other two.

**SMovie 1**. Confocal image for fluorescent *in situ* hybridization. Red = VACHT; Blue = AMPA receptor; Yellow = NMDA receptor.

**SMovie 2**. Confocal image for fluorescent *in situ* hybridization. Green = VGAT; Yellow = NMDA receptor; Magenta = Kainate receptor.
